# *methbiome:* reference-based DNA methylation profiling of long-read metagenomic data

**DOI:** 10.64898/2026.03.28.714822

**Authors:** Ricardo Costeira, Ewan Eustratiou Wong, Jordana T. Bell

## Abstract

**Summary:** Long-read DNA sequencing technologies collect base modification information, but their capacity to profile epigenetic signals has not been widely explored within metagenomic context. Here, we present a novel pipeline consolidating key steps of meta-epigenomic profiling. ***methbiome*** can process both Oxford Nanopore and PacBio long-reads within a reference-based framework, facilitating cross-platform and between-sample comparison of DNA methylation signals in whole microbiomes. The tool is applicable to both microbial communities and isolates, for analyses within both methylation motifs as well as out-of-motif contexts, and can be used in metaepigenome-wide association studies.

**Availability and Implementation:** ***methbiome*** was implemented using *Snakemake* and is available on GitHub (github.com/ricardocosteira/methbiome).

## 1. Introduction

DNA methylation (DNAm) is the main epigenetic mark in bacteria where 6-methyl-adenine (6mA), 5-methyl-cytosine (5mC) and 4-methyl-citosine (4mC) are typically found. Historically challenging to detect, DNAm marks can now be identified at single-base level with long-read (LR) DNA sequencing technologies that profile DNAm on native DNA strands. LR technologies achieve this by detecting modified base-associated changes in enzyme kinetics (PacBio) or electrical current (Oxford Nanopore Technologies, ONT). LR is applied in metagenomics, particularly useful in metagenomic assembly and characterisation of mobile genetic elements, genome structural variation and bacterial strain diversity (1). Despite this, few studies (2–7) have explored DNAm in metagenomic context. Here we present *methbiome* (Figure 1), a novel pipeline that facilitates the analysis of DNAm signals in whole microbiomes. *methbiome* calls DNAm from both PacBio and ONT, generates sample diversity profiles and, crucially, aligns LRs to a reference database while preserving each sample’s DNAm signals, which are included in the mapping output. DNAm scores are estimated at each reference position for different microbes in a sample, allowing for cross-sample analysis of DNAm at the same genetic *loci*.

**Figure 1.**
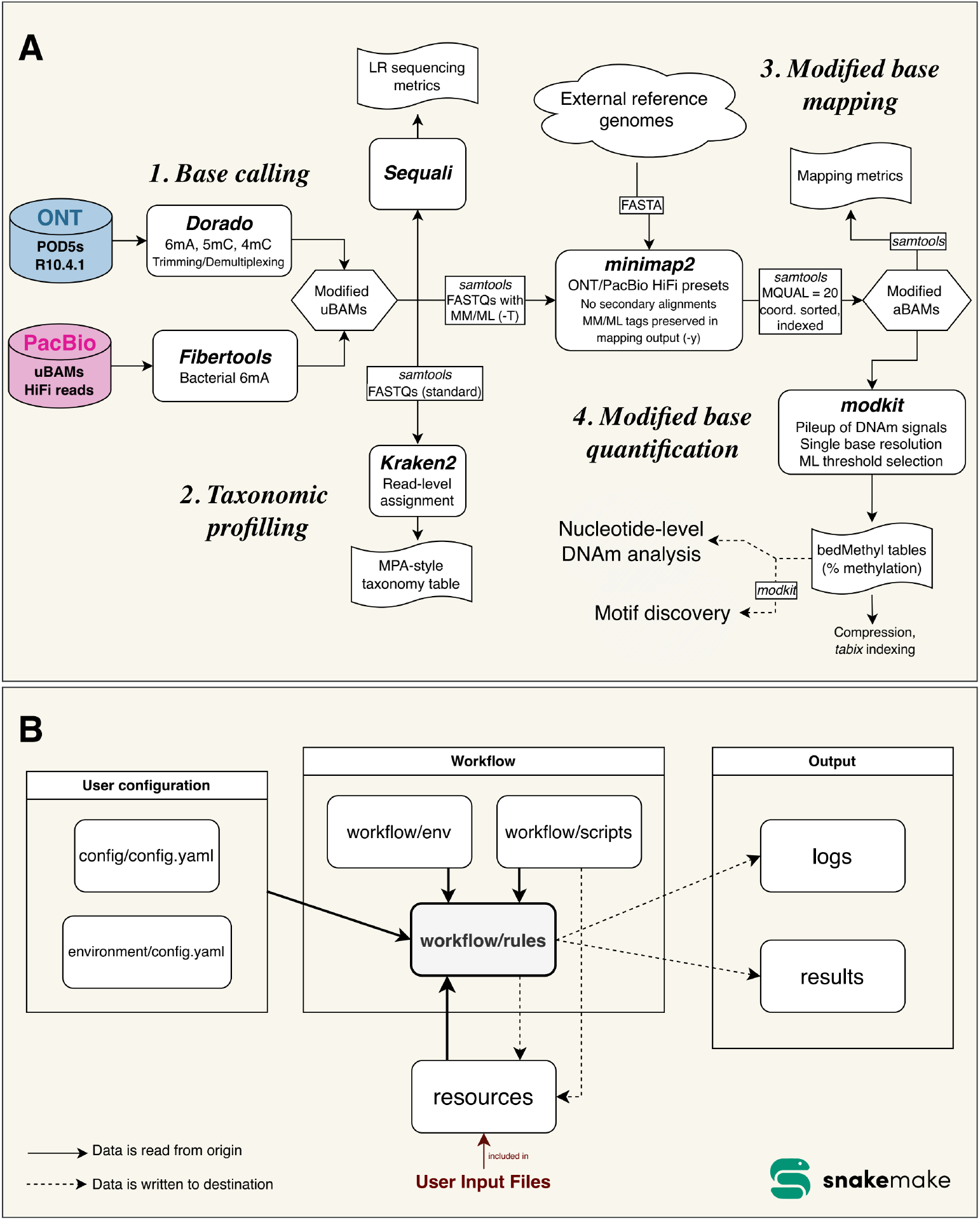
Bioinformatic workflow (A) and software implementation (B) of *methbiome*. Implementation was performed with *Snakemake* where workflow rules orchestrate the interaction between the four modules of the pipeline.

## 2. Bioinformatic workflow

The processing of LRs through *methbiome* can be grouped into four stages: *i)* base calling and data quality assessment; *ii)* taxonomic profiling; *iii)* modified base mapping, and *iv)* modified base quantification (Figure 1A). *methbiome* was implemented using *Snakemake* (8), which manages the flow of information and execution of tasks (Figure 1B). Users can run the pipeline in full or execute it stepwise, completing jobs at designated module checkpoints.

### 2.1. Base calling and data quality assessment

The pipeline input files are user-specified. In the case of PacBio HiFi reads, the pipeline reads in unaligned BAM (uBAM) files from *SMRT Link*. Enzyme kinetics signals from PacBio are first processed with *Fibertools* (9) to call 6mA positions in the long reads. The pipeline does not include a cytosine-specific modified base caller for PacBio at this stage, and *SMRT Link*-processed HiFi reads already include 5mC calls albeit in CpG context. In the case of ONT, the pipeline accepts the location of a *POD5* folder with data from *MinKNOW*. Electrical current signals (squiggles) stored in the *POD5* format from ONT are first processed with Dorado (available at: https://github.com/nanoporetech/dorado) to call 6mA, 5mC and 4mC using an all-context high accuracy model (provided as standard, though configurable by the user). Our implementation of *Dorado* allows the processing of single-plex or multiplexed samples, including steps for trimming and demultiplexing reads from *POD5s*.

Both *Fibertools* and *Dorado* produce uBAM files containing methylation position (MM) and methylation likelihood (ML) data which are standardised SAM/BAM tags encoding DNAm information from sequencing data and are versatile for downstream analyses. The user can choose to name samples at this stage with configuration in *Snakemake*.

After DNAm calling with *Fibertools* and *Dorado*, uBAM files are processed with *Sequali* (10) for quality assessment of the data. *Sequali* produces both visual HTML and text-based reports.

### 2.2. Taxonomic profiling

The uBAM files containing sequence information from module 2.1 are then used to generate taxonomic profiles of the microbiome sample. FASTQ files are generated from the uBAMs with *samtools* (11) for taxonomic assessment with *Kraken2* (12). The current implementation of *Kraken2* uses its prebuilt indexes for speed and compatibility, and automatically downloads the latest “Standard” collection available at https://benlangmead.github.io/aws-indexes/k2 (“Standard”: Refseq archaea, bacteria, viral, plasmid, human and UniVec_Core). Taxonomic assessment in *Kraken2* is performed across all taxonomic levels, including intermediate ranks. Available output formats include the *Kraken* and *MetaPhlAn*-style (mpa-style) format outputs.

Since the *Snakemake* implementation processes one sample at a time, the pipeline includes two post-*methbiome* scripts. The first script allows merging of all mpa-style *Kraken2* outputs into one single taxonomic table using *Kraken2*’s ‘combine_mpa.py’. The second script allows the user to feed various *Sequali* reports into *MultiQC* (13) for easier cross-sample comparison of sequencing metrics. Both scripts are made available in the pipeline GitHub page (github.com/ricardocosteira/methbiome/post-scripts).

### 2.3. Modified base mapping

In the third module of the pipeline DNAm signals stored in the LRs are anchored to a reference database. First, the uBAM outputs from *Fibertools* and *Dorado* are converted to FASTQs using *samtools* which allows the carryover of MM/ML tags to FASTQ headers. Then, the modification-rich FASTQ files are aligned to an external reference with *minimap2* (14) while keeping the MM/ML tags. Mapping is performed using the *minimap2* presets for ONT (“map-ont”) and PacBio HiFi (“map-hifi”) reads according to origin of input data. Secondary alignments and alignments with mapping quality < 20 are excluded from downstream DNAm level calling although another mapping quality threshold (*e*.*g*., 30, 40) can be defined by the user. The resulting aligned BAM (aBAM) file keeps DNAm tags and is coordinate-sorted and indexed against the reference. At this stage alignment metrics are produced using *samtools coverage* and *samtools flagstat* to assess mapping quality and coverage. *methbiome* accepts any microbiome reference in FASTA format. While the pipeline is developed for whole microbiome analysis, it can also analyse DNAm from single isolates with the appropriate genome reference and *Snakemake* configuration (snakemake --profile environment rule_name), where mapping is performed but taxonomic assessment is omitted.

### 2.4 Modified base quantification

DNA methylation scoring is performed on aBAMs from module 2.3 with *modkit* (available at: https://github.com/nanoporetech/modkit). Specifically, the pipeline employs *modkit*’s *pileup* component to aggregate DNAm signals against positions from the whole microbiome reference genomes. A standard ML filter threshold in the pipeline is 0.8 for metaepigenome-wide analysis, but this can be changed in the configuration file. For downstream motif analysis we recommend lower ML scores (*e*.*g*., 65% in line with (15)). For PacBio data *methbiome* implements *modkit*’s *modbam update-tags* to enable cross-platform compatibility with *modkit pileup* in later releases of the software (v. 0.6.0 onwards). Specifically, the pipeline updates PacBio’s aBAM methylation tags to explicitly register ML for all canonical bases in the file so there are no assumptions of methylation status (*e*.*g*., “A+a” → “A+a?” for 6mA). This is the equivalent of using the *--force-allow-implicit* flag in older versions of *modkit pileup*. ONT data remains marked implicit-canonical (*e*.*g*., “A+a.” for 6mA).

Results are tabulated in bedMethyl format where rows correspond to genomic positions, taking strand orientation into account. The table includes several metrics, with the key ones being the percentage of modification and coverage at each base. Because whole microbiome DNAm tables can be large, we implemented the option to compress and index the bedMethyl file with *bgzip* and *tabix* (16). Selecting this option can be useful in downstream analyses that aim to query specific microbes or regions of interest, based on genome reference headers and coordinates, with *samtools* in Bash or *Rsamtools* combined with the *GenomicRanges* package (17) in *R*.

The outputs of *methbiome* support association analyses at single-nucleotide base resolution. Downstream motif-level characterization of DNAm can also be performed using the bedMethyl tables generated and other *modkit* modules, with *modkit search* enabling *de novo* motif discovery and *modkit evaluate* supporting the interrogation of motifs defined *a priori*. Integration of *tabix* in *methbiome* allows the query of target motifs in individual bacterial genomes by specifying the *--contig* flag and relevant reference headers in downstream *modkit* searches.

Future versions of *methbiome* aim to incorporate motif discovery as well as provide the option to remove human reads from LRs in studies requiring host decontamination prior to DNAm quantification.

## 3. Software implementation

*Snakemake* manages data transfer between different components of *methbiome* including user configurations, workflows, resources and output (Figure 1B). The configuration files guide the workflow’s execution by providing module specific parameters (*i*.*e*., execution parameters of software packages) and environment settings (*i*.*e*., resources allocated to individual jobs). The user must edit the configuration files (“config/config.yaml” and “environment/config.yaml”) to run the pipeline. The user configuration is read by the workflow component which manages the execution. The workflow contains the sub-components *i) rules*, which manage the logic that dictates the flow of processes and orchestrate the interaction between all components of the pipeline (represented in Figure 1A); *ii) env*, which holds definitions of the conda environment built by *Snakemake* (*i*.*e*., self-contained, isolated spaces with specific versions of software packages) and is accessed by *rules* to execute each step of the pipeline; and *iii) scripts*, which stores external helper scripts that are executed by *rules*. Resources are files that are either retrieved or provided by the user. These include databases or software packages that cannot be defined in conda environments as well as input files from the user. The pipeline generates two key outputs: *i) results* which are the outcomes of the workflow, and *ii) logs* which stores execution logs of each step of the pipeline and reports failed jobs submitted to the High-Performance Computing (HPC) server.

## 4. Application

The pipeline was applied to publicly available data releases from ONT and PacBio for the microbiome standard *ZymoBIOMICS Fecal Reference with TruMatrix Technology* (Zymo Research, D6323). The D6323 sample was sequenced by ONT on two PromethION flow cells using R10.4.1 chemistry (available at: https://epi2me.nanoporetech.com/zymo_fecal_2025.05/) and by PacBio on two Revio SMRT Cell systems (available at: https://www.pacb.com/connect/datasets/). Using *methbiome*, data from the two ONT replicates (yielding 184.78 and 91.32 gigabases after base call) and PacBio replicates (yielding 67.98 and 67.08 gigabases) were processed with maximum runtimes of 96.2 hours and 15.9 hours, respectively (Supplementary Table 1). Analyses were conducted in a HPC environment with request of 60 CPU cores, one NVIDIA GPU, and 128 GB of RAM. The HPC system was running Ubuntu 22.04.5 LTS on an x86_64 architecture with Slurm (v. 25.05.5). *methbiome* was run with *Snakemake* v. 9.8.0 in a *tmux session (tmux* v. 3.5). Dorado base calling and LR mapping were the most time-constraining steps of the pipeline (Supplementary Table 1). In this application, we mapped LRs to a 2.7 GB reference FASTA file containing only highly important strains from *proGenomes4* (18). Employment of non-redundant context-guided databases is recommended for speed and coverage.

## 5. Conclusion

*methbiome* is a novel pipeline to process raw base modification signals from PacBio and ONT long-reads for epigenetic profiling and contextualise these within a metagenomic framework. The reference-based approach provides a stable and accurate coordinate system essential for robust epigenetic analysis, enabling rapid and reliable annotation of DNAm signals to motifs and nearby genes in downstream analyses. Moreover, by anchoring DNAm signals to a reference, the pipeline allows consistent comparison of epigenetic patterns across loci, samples, and studies, while substantially reducing the computational burden associated with constructing metagenome-assembled genomes (MAGs). Finally, results from *methbiome* can be seamlessly integrated into future metaepigenome-wide association studies, enabling comprehensive analysis of DNAm signals across diverse microbiomes and their environmental contexts.

## Supplementary information

**Supplementary Table 1.** Resources used by *methbiome* in HPC environment.

| Rule | Name of software package | Version | Number of processing units | Memory limit | Runtime (dd:hh:mm:ss) |  |  |  |
| --- | --- | --- | --- | --- | --- | --- | --- | --- |
|  |  |  |  |  | ONT ZymoBIOMICS D6323 (PAU85136) 2.2 TB: 184.78 Gbs and 16.3 M reads | ONT ZymoBIOMICS D6323 (PAW10254) 1.2 TB: 91.32 Gbs and 10.7 M reads | PacBio ZymoBIOMICS D6323 (224423_s1) 33 GB: 67.98 Gbs and 8.9 M reads | PacBio ZymoBIOMICS D6323 (231547_s2) 32 GB: 67.01 Gbs and 8.7 M reads |
| *install_dorado* | python | 3.14.2 | 1 CPU | 128 GB | 00:00:01:05 | 00:00:01:53 | -- | -- |
|  | beautifulsoup4 | 4.14.2 |  |  |  |  |  |  |
|  | requests | 2.32.5 |  |  |  |  |  |  |
| *install_kraken2_db* | python | 3.14.2 | 1 CPU |  | 00:00:46:38 | 00:00:52:51 | 00:00:43:49 | 00:00:46:51 |
|  | beautifulsoup4 | 4.14.2 |  |  |  |  |  |  |
|  | requests | 2.32.5 |  |  |  |  |  |  |
| *dorado* | dorado | 1.3.1 | 1 GPU |  | 01:07:05:25 | 01:00:23:23 | -- | -- |
| *fibertools_predict* | fibertools-rs | 0.8.1 | 60 CPU |  | -- | -- | 00:01:41:17 | 00:01:38:23 |
| *sequali* | sequali | 1.0.2 | 60 CPU |  | 00:00:55:51 | 00:00:26:34 | 00:00:09:10 | 00:00:09:06 |
| *kraken2* | kraken2 | 2.17.1 | 60 CPU |  | 00:07:39:37 | 00:04:51:18 | 00:02:59:36 | 00:03:04:36 |
|  | krakentools | 1.2.1 | 1 CPU |  |  |  |  |  |
| *bam_to_fastq* | samtools | 1.23 | 60 CPU |  | 00:14:32:09 | 00:06:31:39 | 00:04:35:03 | 00:04:22:23 |
| *minimap2\** | minimap2 | 2.3 | 60 CPU |  | 01:09:25:29 | 00:14:01:34 | 00:04:16:31 | 00:04:17:54 |
|  | samtools | 1.23 |  |  |  |  |  |  |
| *filter_samtools* | samtools | 1.23 | 60 CPU |  | 00:05:36:21 | 00:02:38:54 | 00:00:52:43 | 00:00:51:55 |
| *modkit* | ont-modkit | 0.6.1 | 60 CPU |  | 00:02:02:52 | 00:01:04:46 | 00:00:38:59 | 00:00:38:43 |
|  | samtools | 1.23 |  |  |  |  |  |  |
| *tabix* | tabix | 1.11 | 60 CPU |  | 00:00:07:55 | 00:00:06:03 | 00:00:00:53 | 00:00:00:28 |
|  | samtools | 1.23 |  |  |  |  |  |  |

\*Mapping was performed using the representative genomes of highly important strains contained in *proGenomes4*. We mapped ONT and PacBio samples against the combined genomes of 751 microbial strains (genomes file: https://progenomes.embl.de/data/repGenomes/pg4_genomes_representatives.fna.gz; list of highly important strains used to subset genomes file: https://progenomes.embl.de/data/pg4_highly_important_strains.tsv.gz).

## Acknowledgements

We acknowledge use of King’s Computational Research, Engineering and Technology Environment (CREATE, https://doi.org/10.18742/rnvf-m076) in this work.

## Author contributions

RC and JTB supervised the study. RC conceptualised and wrote the pipeline. JTB oversaw the study and secured its funding. EEW implemented the pipeline in *Snakemake*. RC, EEW and JTB wrote, reviewed and approved the manuscript.

